# Genomic and phenotypic characterization of novel jumbo and small bacteriophages infecting *Xanthomonas hortorum* pv. *vitians*: Toward a phage-based biocontrol strategy

**DOI:** 10.1101/2025.08.08.669339

**Authors:** Anaelle Baud, Inès Rougis, Nicolas Taveau, Denis Costechareyre, Marie Graindorge Beaume, Franck Bertolla

## Abstract

In recent years, the use of bacteriophages as biocontrol agents has gained renewed attention, especially for targeting major phytopathogenic genera like *Xanthomonas*. However, their potential against *Xanthomonas hortorum* pv. *vitians*, the causal agent of bacterial leaf spot of lettuce, remains largely unexplored with only two lytic phages reported to date. In this study, eight novel lytic tailed phages were isolated from wastewater samples, including three jumbophages, which are the first known to infect this pathovar. These jumbophages define a new viral genus and exhibit distinct biological characteristics, including broader host ranges, but lower burst sizes and slower lytic cycles compared to the smaller-genome phages. Together, the newly isolated phages exhibited lytic activity against nearly all *X*. *hortorum* pv. *vitians* strains tested while remaining highly specific to the pathovar. Genomic analyses revealed strictly lytic profiles, and absence of virulence or resistance genes, supporting their biosafety for agricultural use. Furthermore, the isolation of a phage infecting a bacterial mutant previously shown to be resistant to the virulent phage Φ*Xhv*-1 highlights the potential for designing robust phage cocktails to mitigate resistance emergence. All phages demonstrated high stability across a wide range of pH and temperature conditions, but remained highly sensitive to UV exposure. These findings provide a valuable foundation for developing phage-based biocontrol strategies against bacterial leaf spot of lettuce.

**IMPORTANCE:** Ranked among the leading plant-pathogenic bacterial genera, *Xanthomonas* species affect a wide range of crops and contribute to major agricultural losses worldwide. Among these, *Xanthomonas hortorum* pv. *vitians*, the causal agent of bacterial leaf spot of lettuce, poses a serious threat to lettuce production worldwide. Currently, no effective disease managements strategies are available to reduce this foliar hemibiotropic phytopathogen. Bacteriophage-based biocontrol has shown promising results against *Xanthomonas* spp. However, its application against this pathogen remains largely unexplored, with only two phages isolated to date. Successful phage application in agriculture requires careful selection and through characterization to ensure safety and efficacy. In this study, the phage diversity targeting this phytopathogen was expanded through the isolation of eight novel lytic phages, including the first jumbophages infecting this species. Their genomic and biological properties provide a valuable foundation for developing future phage-based treatment against this challenging pathogen.

## INTRODUCTION

Ensuring global food security in light of the projected population growth to 9.6 billion by 2050 will require an estimated 70-80% increase in agricultural production, according to the Food and Agricultural Organization of the United Nations (FAO). However, plant pathogens are already responsible for up to 41% yield losses in major crops, resulting in an estimated annual economic impact of US$220-290 billion and affecting approximately 10% of the global food supply (2, 3). To date, more than two hundred bacterial plant pathogens have been identified. Reported yield losses caused by these bacteria vary widely depending on the crop and region, ranging from 10% to 40% (4, 5). Climate changes, particularly rising temperatures, are expected to exacerbate this burden by altering pathogen life cycles, expanding their geographic range, and increasing the severity of plant-pathogen interactions (6, 7). In this context, biocontrol strategies using microbial agents are emerging as promising and sustainable alternatives to chemical pesticides, especially copper-based compounds, whose persistence in soil negative impact on beneficial microbiota, and contribution to bacterial resistance raise significant concerns (8, 9). The development and implementation of these environmentally friendly approaches are increasingly supported by public policies, such as the European Green Deal, which aims to reduce synthetic pesticide use by 50% by 2030. Faced with these new challenges, growing attention has been directed toward bacteriophages, viruses that specifically infect bacteria. First discovered by Frederick Twort and Félix d’Herelle in the early 20th century, phages have a long-standing history in biocontrol applications (10, 11). Their potential was recognized as early as 1924, when Hemstreet and Mallmann used a filtrate from decomposed cabbage containing phages to treat *Xanthomonas campestris* pv. *campestris* (12). The first documented success in field application reportedly occurred in 1935, when phage treatment was used to control Stewart’s wilt diseases, caused by *Pantoea stewartia* (13).

Despite this early interest, phage-based biocontrol has since remained marginal for decades due to technical limitations, regulatory gaps, and a lack of genomic resources. However, recent advances in phage genomics, combined with the growing demand for sustainable agriculture, have renewed scientific and industrial interest in isolating and characterizing novel phages targeting key pathogens. Indeed, phages present unique biological features for biocontrol applications: (i) high specificity toward bacterial targets, (ii) safety for eukaryotic cells, (iii) natural abundance and genetic diversity, enabling the identification of specific phages for virtually any bacterial host, and (iv) ability to multiply exponentially within bacteria through successive lytic cycles, making them potentially cost-effective to produce and apply (3, 14, 15). Since 2005, several private companies (e.g., OmniLytics Inc., Enviroinvest Corp., A&P Inphatec, APS Biocontrol Ltd.) have developed and commercialized phage-based biopesticides for both preventive and post-harvest applications targeting major phytopathogenic bacteria. Among these pathogens, *Xanthomonas* ranks among the ten most important phytopathogenic bacterial genera, causing diseases in over 400 plant species, including economically critical crops such as rice, tomato, and cabbage (8, 16, 17). Among its pathovars, *X*. *hortorum* pv. *vitians* is the causal agent of bacterial leaf spot of lettuce (*Lactuca sativa*, *Asteraceae*) (18). This bacteriosis is characterized by the development of water-soaked lesions on leaves, which expand and coalesce, ultimately leading to the necrosis of entire leave (19). This phytopathogen is present worldwide and infects all cultivated types of lettuce (20).

Symptomatic lettuces are often unmarketable, and under favorable conditions, particularly cool and wet weather, several outbreaks have been reported to affect up to 100% of field-grown lettuce. In the absence of effective preventive or curative measures, this disease remains a serious threat to lettuce producers, causing substantial yield and economic losses (21). Current management practices rely mainly on prophylactic measures such as crop rotation, cultivar selection, and sanitation, underscoring the urgent need for alternative solutions (22, 23). Recent studies have demonstrated the biocontrol efficacy of phages against *Xanthomonas* spp. phylogenetically related to *X*. *hortorum* pv. *vitians* under both greenhouse and field conditions. For example, a phage cocktail (FoX2-FoX6) reduced the proportion of plants infected by *X*. *campestris* pv. *campestris* by up to 30% in field trials (24). Against *X*. *oryzae* pv. *oryzae*, phage Xoo-sp2 reduced disease symptoms by one-third (25) while phage X3 achieved a reduction of 83.1% (26). Additional successful phage applications are reviewed by Holtappels *et al*. (8). In contrast, phages active against *X*. *hortorum* pv. *vitians* remains largely overlooked, with only two lytic phages described to date (27). This marked underrepresentation highlights a broader gap in phage research for this pathovar. Expanding this limited phage collection and thoroughly characterizing their biological and genomic properties are crucial steps toward the development of a robust and sustainable phage-based biocontrol solution.

In this study, we report the isolation of eight novel bacteriophages targeting *X. hortorum* pv. *vitians*, including 3 jumbophages which are the first known to infect this pathovar. To assess their potential for biocontrol application, we performed phenotypic and genomic characterization. This included host range profiling across the genetic diversity of the pathovar *vitians* and other *Xanthomonas*, bacterial growth inhibition assays, adsorption kinetics, and one-step growth curve analysis. In addition, their environmental stability under key abiotic stresses, including pH, temperature, and UV exposure was also evaluated. Finally, whole-genome sequencing was used to assess their genetic features, taxonomic classification, and biosafety compliance criteria relevant to agricultural applications.

## MATERIALS AND METHODS

### Strains and growth conditions

*Xanthomonas* strains used in this study are listed in **Table S1**. Strains were routinely streaked onto tryptic soy agar (TSA) plates and incubated at 28°C for 48-72 h. Single colonies were used to inoculate tryptic soy broth (TSB) and cultured overnight at 28°C with shaking at 160 rpm. Strains of *X. hortorum* pv. *vitians* used for phage isolation were listed in **Table S1**. Semi-solid agar overlays consisting of TSB with 0.6% agar (or 0.4% for jumbophages), supplemented with 1 mM CaCl_2_ to promote phage adsorption, were used for plaque assays. For phage infection experiments, overnight bacterial cultures were diluted in TSB supplemented with 10 mM CaCl_2_ to an optical density at 600 nm (OD_600_) of 0.15 and incubated at 28°C with shaking at 160 rpm for 1 hour to reach the early logarithmic phase prior to phage addition. All strains were kept at −80 °C in 30% (v/v) glycerol vials for long-term storage.

### Phage isolation, purification, and propagation

A total of 42 samples were collected from various wastewater treatment plants in France for the isolation of *X*. *hortorum* pv. *vitians* phages. Phages were isolated using an enrichment-based approach and a screen by the double-layer agar technique as adapted from Plumet *et al*. (28). Filtered effluent were mixed with early logarithmic-phase cultures of different *X. hortorum* pv*. vitians* strains and reprocessed by centrifugation and filtration steps after an overnight incubation. The presence of phage was evaluated using the double-layer agar technique, where the appearance of individual plaques indicated successful phage isolation. Individual plaques were purified through two successive re-isolation steps. After incubation at 4°C for 1 h and filtration through a 0.2 µm filter, phage stocks were stored at 4°C in SM buffer. Plaque morphology (i.e., size, clarity, turbidity, edge definition) was evaluated using the single agar layer method. Unless otherwise specified, all phage suspensions used in the following experiments were filtered through 0.2 µm pore-size filters.

Phages were propagated using the double-layer agar technique. Plates with confluent lysis were flooded with SM buffer, and the resuspended soft top agar were collected and centrifuged 10 min at 5,000 x *g* at 4°C. Supernatants were filtered through a 0.2 µm filter before storage at 4°C.

For large-scale production, each phage was amplified in liquid bacterial culture. An early logarithmic-phase culture of the production strain was mixed with phage at a multiplicity of infection (MOI) of 0.1 and incubated for up to 16 h at 28°C. Following lysis, cultures were centrifuged and the clear supernatants were filtered. Phage titers (in PFU/mL) were determined by the spot assay method on double-layer agar.

### Phage morphological characterization by transmission electronic microscopy (TEM)

The preparation of phages samples for phage samples for TEM observation was performed following the protocol previously described by Plumet *et al*. (28), with minor modifications. Briefly, phage particles were concentrated by centrifugation and washed twice in acetate ammonium buffer (NH4-acetate 0.1M). Grids were negatively stained using 2% uranyl acetate and visualized with a JEOL 1400 Flash transmission electron microscope operated at 120 kV. Virion dimensions (capsid diameter, tail length/width) were measured from at least ten individual virion micrographs using ImageJ software (v.154d, (29)). Results are reported as a mean value ± standard error.

### Phage whole-genome sequencing and analysis

Phage DNA was extracted and purified as described previously by Gendre J (30) without addition of SDS and proteinase K. Libraries were prepared using the Illumina TruSeq PCR-Free and Nextera XT DNA Library Prep Kits, and sequencing was performed on an Illumina Miseq or NovaSeq 6000 S4 platforms (paired-end, 150 bp), depending on the phage sample.

Reads quality was assessed using FastQC v0.12.1 (31) followed by a cleaning using PRINSEQ 0.20.4 (32). Reads were assembled using SPAdes 4.0 (33). Contigs sequences were queried against the NCBI nucleotide (nt) database using BLASTn (34) to confirm viral origin and exclude contaminant sequences. Assembly quality was evaluated by read mapping using Bowtie2 v2.5.4 (35), followed by processing with Samtools v1.20 (36). Structural and functional annotation was carried out using the rTOOLS pipeline (RIME Bioinformatics). Transfer RNAs were identified using tRNA scan-SE v2.07 (37). Phage lifestyle was predicted using PhageAI (38). Comparative genomic analysis among phage isolates was performed using CLINKER (39). Phylogenomic relationships were inferred with ViPTree (40), which constructs a proteomic tree based on genome-wide tBLASTx similarities against reference viral genomes. Intergenomic similarities among complete phage genomes were calculated using VIRIDIC with blastn default settings (41).

### Host range determination

The lytic host range of the phages was assessed by spot assays on 41 *Xanthomonas* strains, including 34 *X*. *hortorum* pv. *vitians*, 5 strains from other pathovars and 2 from other species (**Table S1**). Briefly, 10-fold serially diluted phage suspensions were spotted onto log-phase bacterial cultures in soft agar overlays and incubated overnight at 28°C. A strain was considered susceptible if clear countable plaques were observed upon phage application. Lysis from without (halo without plaques) or absence of lysis indicated resistance. Efficiency of plating (EOP) was calculated as the ratio of the phage titer (in PFU/mL) on the test strain to the titer on the original isolation host. Strains were classified as highly sensitive (EOP > 0.5), sensitive (10 ^-4^ < EOP < 0.5) or resistant (EOP < 10 ^-4^). This classification was adapted from previously established EOP categories (42, 43), using a lower resistance threshold (EOP < 10^-4^ instead of 10^-3^). All experiments were done in duplicate, with two technical replicates per phage dilution. The relationship between phage -genome size (categorized as small or large) and host range breadth was analyzed using a chi-squared test of independence. To evaluate whether genetic diversity among *X*. *hortorum* pv. *vitians* strains influenced phage susceptibility, differences in overall susceptibility scores across MLSA-defined groups (A, B, and C) were tested using the Kruskal-Wallis rank-sum test. Group-specific biases in susceptibility at the individual phage level were assessed using Fisher’s exact tests with Monte Carlo simulation (10,000 replicates, to approximate p-values), followed by Bonferroni correction for multiple testing.

### Bacterial growth inhibition assay

The ability of phages to inhibit the growth of their production strain was assessed by monitoring OD_600_ using a Bioscreen C MBR BACTERIO (Thermo Fisher Scientific). Briefly, early exponential-phase host cultures were inoculated with phage suspensions at MOIs ranging from 0.00001 to 50 in 100-well Honeycomb plates. After a 30 min adsorption period at room temperature, OD_600_ was recorded every 20 min for 24 h at 28°C under continuous orbital shaking. TSB only and the bacterial strain without phages served as negative and bacterial growth controls, respectively. Four technical replicates per treatment were performed.

### Adsorption assays

Phage adsorption kinetics were determined by quantifying non-adsorbed phages over time. Early-log-phase bacterial cultures were mixed with phages at a MOI of 0.01. Co-cultures were incubated at 28°C without agitation, except for jumbophages, which were incubated with gentle shaking at 50 rpm. Aliquots were collected at defined intervals, filtered through 0.2 µm pore-size filters to separate free phages, and then quantified by spot assay. The percentage of free phages was calculated relative to the initial titer. The adsorption rate constant (*k*) was calculated as described previously (44). Two independent experiments were performed, each comprising three technical replicates.

### One-step growth curves

One-step growth curves were performed to assess phage infectivity and replication dynamics. Phages were added at a MOI of 0.1 to early log-phase bacterial cultures and incubated at 28°C (with gentle agitation for jumbophages) for a pre-determined adsorption time. Non-adsorbed phages were removed by centrifugation (5,000 x *g*, 21°C, 10 min), and the cell pellets were washed twice in 35 mL of TSB supplemented with 10mM CaCl_2_. Resuspended co-cultures were incubated at 28°C with shaking at 160 rpm, and aliquots were collected at various time points for free phage quantification by spot assay. Burst size was calculated as the ratio of phages released after the rise period to the number of adsorbed phages. Two independent experiments were performed, each with three technical replicates per time point.

### Effect of temperature, pH, and ultraviolet irradiation on phages stability

To assess phage stability, suspensions were subjected to various abiotic conditions including temperature, pH, and UV-B irradiation. For thermal stability, phages were incubated in the dark for 1 h at temperatures ranging from 4°C to 75°C in a thermal cycler (Analytik Jena). pH stability was evaluated by incubating phages in SM buffer adjusted to pH values from 2 to 12 at room temperature, in the dark, for 1 h. For UV sensitivity assays, phage suspensions were poured into sterile empty Petri dishes, left uncovered and exposed to UV-B light emitted by a Bio-Link Crosslinker BLX-E312 (Vilber Lourmat) at a 16 cm distance for 0, 2, 5, 10, 15 or 30 min. Following each treatment, phage viability was assessed by plaque counting using the double-layer agar method. The phage survival rate (%) was calculated relative to the initial titer measured under optimal storage conditions (4°C, pH 7.5, in the dark). Each abiotic condition was tested in two independent experiments, each with three technical replicates.

### Data analysis and visualization

All figures were generated using R (v4.3.3; R Core Team) within RStudio (version 2023.12.1.402 “Ocean Storm” Release; RStudio Team), unless otherwise specified. Visualizations were primarily created using the ggplot2 package (v3.5.1), with additional packages including readxl (v1.4.3) for data import, dplyr (v1.1.4) for data manipulation, and patchwork (v1.3.0) for figure assembly. Specific visualizations, such as heatmaps, employed ComplexHeatmap (v2.18.0) and circlize (version 0.4.16). Other packages used for figure-specific processing include scales (v1.3.0), tidyr (v1.3.1), purr (v1.0.2), cowplot (v1.1.3), and viridis (v0.6.5). When specified, figures were post-processed for layout optimization and labeling using Inkscape (v1.4). Figures were exported in high resolution (1200 dpi) in SVG, PNG, and/or PDF formats for publication purposes.

Additional methodological details are provided in the **Supplementary Methods**.

## RESULTS

### Eight phages isolated from wastewater show distinct plaque morphologies and *myovirus*-like structures

Eight novel phages infecting *X. hortorum* pv. *vitians* were isolated from wastewater samples. The phages Φ*Xhv*-2, Φ*Xhv*-5, Φ*Xhv*-7, Φ*Xhv*-12, Φ*Xhv*-15, Φ*Xhv*-16, Φ*Xhv*-18, and Φ*Xhv*-28 were isolated on different bacterial strains representative of the pathovar’s genetic diversity, complemented by a LPS mutant lacking the side branches of the O antigen (i.e., AB16734 ΔLPS3) (**Table S1**). All phages formed clear plaques on their respective production strains, except for Φ*Xhv*-16, which produced slightly turbid plaques **(Fig. 1A)**. Plaque morphology and diameter are summarized in **Table S2** and ranged from 0.563 ± 0.012 mm (Φ*Xhv*-15) to 2.392 ± 0.561 mm (Φ*Xhv*-7). Notably, phages Φ*Xhv*-12, Φ*Xhv*-15, and Φ*Xhv*-16 formed smaller plaques, with a mean diameter of 0.68 mm. In contrast, the five other phages produced larger plaques averaging 1.93 mm, corresponding to approximately a 2.8-fold size difference. TEM micrographs analysis revealed that all eight phages had a *myovirus*-like morphology, characterized by an icosahedral capsid and a contractile tail (**Fig. 1B**). Capsid diameters of phages Φ*Xhv*-12, Φ*Xhv*-15, and Φ*Xhv*-16 (**Table S2**) exceeded 110 nm, making them almost twice as large as those of the other five phages, which ranged from 61.1 ± 1.3 nm (Φ*Xhv*-28) to 73.6 ± 1.3 nm (Φ*Xhv*-7). Tail lengths ranged from 86.3 ± 7.3 nm to 118.6 ± 1.5 nm, with no major differences observed across the phage set.

**Figure 1.**
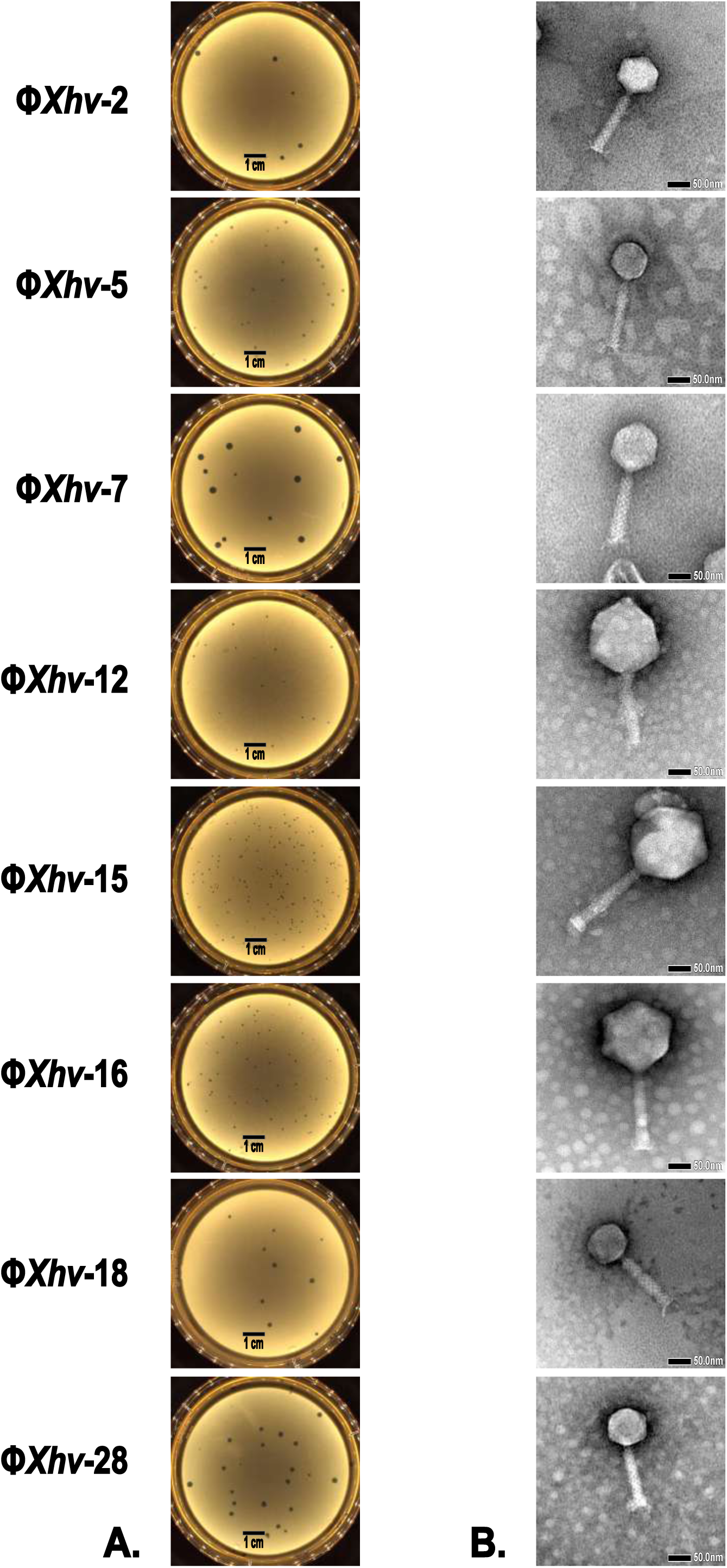
Morphological characterization of novel bacteriophages infecting *X. hortorum* pv. *vitians*. **A.** Plaque morphology of eight distinct bacteriophages (Φ*Xhv*-2, Φ*Xhv*-5, Φ*Xhv*-7, Φ*Xhv*-12, Φ*Xhv*-15, Φ*Xhv*-16, Φ*Xhv*-18, and Φ*Xhv*-28) isolated from sewage samples. Phages were propagated on lawns of *X*. *hortorum* pv. *vitians* strains. Images show representative plaque morphologies observed after overnight incubation at 28°C on tryptic soy agar (TSA). Black scale bars = 1 cm. **B.** Transmission electron micrographs (TEM) showing the virion morphology of the corresponding phages, negatively stained with uranyl acetate. All phages display the typical *myovirus* morphotype. Black scale bars = 50 nm.

### Broad host range and distinct infectivity profiles highlight two major phage groups

The host range of these eight phages was assessed using a panel of 34 *X*. *hortorum* pv. *vitians* isolates and 7 strains from other pathovars or species of *Xanthomonas* (**Fig. 2**). Together, the phages lysed 94% (32 out of 34) of the *vitians* isolates. Only two strains (LM17692 and LM17388) were resistant to all tested phages. Based on EOP values, the phages grouped into three distinct infectivity profiles. Φ*Xhv*-12, Φ*Xhv*-15, and Φ*Xhv*-16 exhibited the broadest host ranges, each infecting more than 25 *X*. *hortorum* pv. *vitians* isolates with moderate to high EOP values (> 10^-3^). This broad host range was significantly associated with their larger genome sizes, as confirmed by a chi-squared test (χ^2^ = 84.294, df = 1, *p.value* = 2.2 × 10^-16^). In contrast, Φ*Xhv*-2, Φ*Xhv*-5, Φ*Xhv*-7, and Φ*Xhv*-18 displayed narrower host ranges, infecting between 3 (Φ*Xhv*-2) and 12 (Φ*Xhv*-5) isolates, corresponding to 8% to 35% of the tested panel. A third highly specific profile was represented by Φ*Xhv*-28, which exhibited lytic activity exclusively against the LPS3 mutant strain (AB16734 ΔLPS3). The potential influence of *X*. *hortorum* pv. *vitians* genetic diversity on phage susceptibility was evaluated by comparing sensitivity across MLSA-defined groups. A first analysis based on overall phage susceptibility scores revealed no significant difference between groups A, B, and C (H = 3.36, df = 2, *p.value* = 0.186), suggesting a similar average response to the phage panel across the genetic groups. To explore potential group-specific interactions at the individual phage level, each phage was then analyzed individually. Among the eight phages tested, only Φ*Xhv*-5 displayed a statistically significant bias, preferentially infecting strains from group C (adjusted *p. value* = 0.02 after Bonferroni correction). Due to limited representation in some categories, potential effects of strain isolation year and geographic origin could not be assessed. None of the phages exhibited lytic activity against non-*vitians* strains supporting their high level of pathovar specificity. This nearly complete lytic activity within *X*. *hortorum* pv. *vitians*, combined with the absence of off-target effects, underscores the potential of these phages for targeted biocontrol applications.

**Figure 2.**
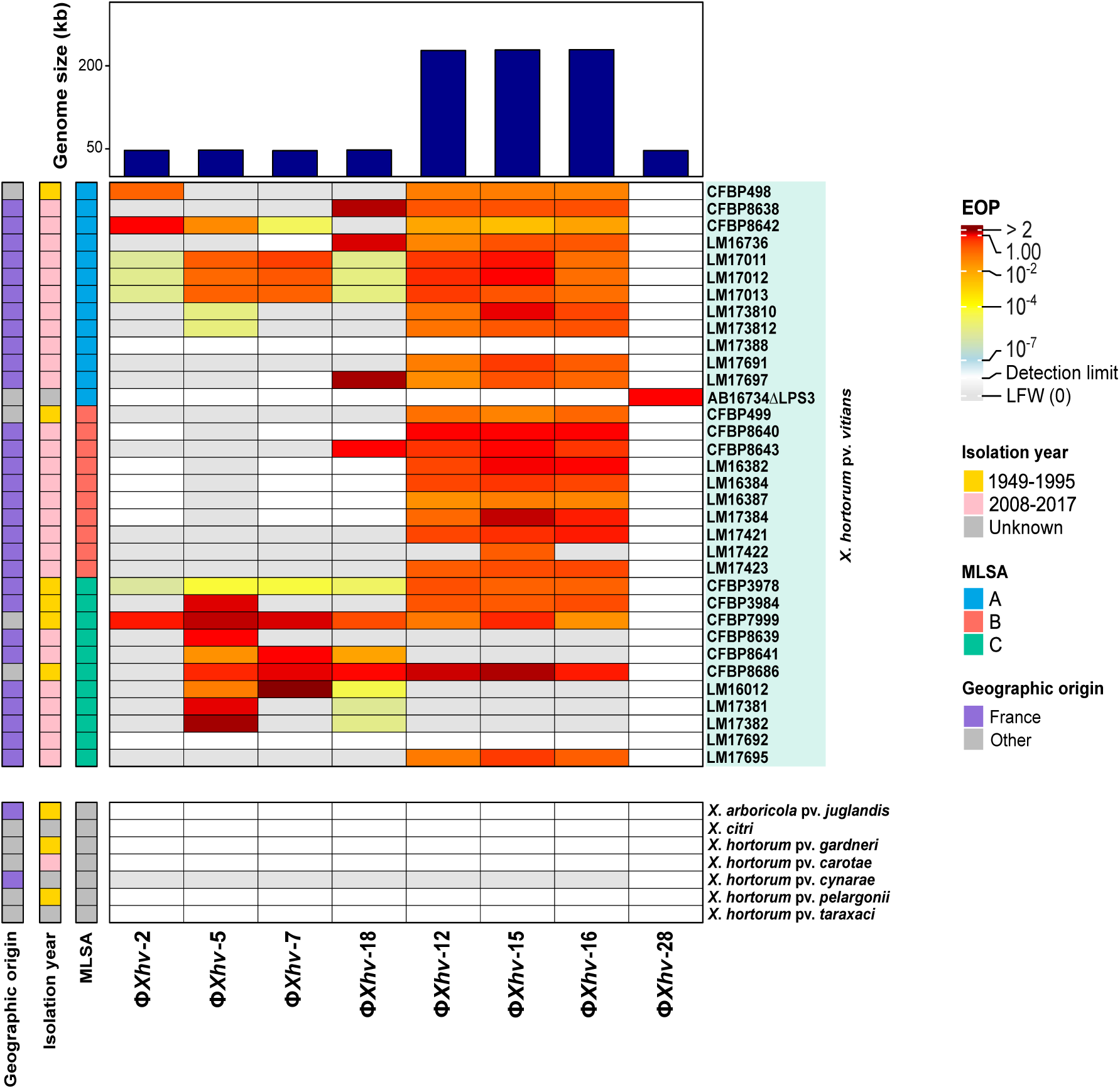
Host range and efficiency of plating of novel phages active against *X. hortorum* pv. *vitians*. The heatmap shows the efficiency of plating (EOP) of eight phages (Φ*Xhv*-2, Φ*Xhv*-5, Φ*Xhv*-7, Φ*Xhv*-12, Φ*Xhv*-15, Φ*Xhv*-16, Φ*Xhv*-18, and Φ*Xhv*-28) tested against a panel of 41 *Xanthomonas* strains, comprising 34 strains of *X. hortorum* pv. *vitians* and 7 strains of other *Xanthomonas* species (bottom rows). EOP values are represented by a color scare ranging from red (> 1) to light yellow (10^-4^) for high to moderate levels of infection. Gray indicates resistant strains or no lysis (including lysis from without, LFW) and white color indicates the detection limit. The bar chart above the heatmap depicts the genome size (in kb) of each phage, as determined by genome sequencing. The left-side annotation columns provide metadata for each bacterial strain, including multilocus sequence analysis (MLSA) group (A, B, or C) (18), year of isolation (1949-1995 in yellow, 2008-2017 in red, unknown in gray), and geographic origin (France or other). All experiments were performed in two independent biological replicates, each with two technical replicates, and representative data are shown.

### Phages infection kinetics reveal differences in adsorption and lytic efficiency

Growth inhibition assays confirmed the lytic nature of the eight phages on their respective isolation strains across a range of MOIs (**Fig. S1**). For all newly isolated phages, bacterial growth inhibition was dependent on phage concentration. Φ*Xhv*-5 and Φ*Xhv*-7 exhibited the fastest lytic activity, fully suppressing bacterial growth within 3 h even at low MOI (i.e., 0.01), demonstrating the high efficiency of their lytic cycle. Even lower MOIs (down to 0.0001) achieved complete inhibition of growth in over 5 h without any regrowth. For Φ*Xhv*-2, Φ*Xhv*-18, and Φ*Xhv*-28, a higher MOI (≥ 1) was required to rapidly lyse the entire bacterial population. Lower MOI (< 0.1) were insufficient to fully inhibit bacterial growth within 10 h for Φ*Xhv*-18 and Φ*Xhv*-28. In contrast, phages Φ*Xhv*-12, Φ*Xhv*-15 and Φ*Xhv*-16 exhibited a much slower lytic cycle. Bacterial growth was observed for more than 2 h even at high MOIs (> 1), with inhibition occurring only after more than 5 h at MOIs above 0.1. To gain further insights into the phage infection process, adsorption efficiency and kinetics parameters were measured. Adsorption efficiency varied among the phages. Phages Φ*Xhv*-2, Φ*Xhv*-5, and Φ*Xhv*-7 exhibited rapid adsorption, with over 90% of virions adsorbed within 20 min **(Fig 3.A)**. Adsorption was slightly slower for Φ*Xhv*-18 and Φ*Xhv*-28, which required 30 to 40 min to reach similar levels. In contrast, Φ*Xhv*-12, Φ*Xhv*-15, and Φ*Xhv*-16 displayed slower and incomplete adsorption dynamics **(Fig3.B)**. For Φ*Xhv*-12, only ∼45% of virions were adsorbed after 30 min of incubation. Φ*Xhv*-15, and Φ*Xhv*-16 were the slowest, with more than one-third of the phage population remaining unabsorbed even after 60 to 90 min of incubation. Adsorption constants, kinetic parameters, and burst sizes, calculated from one-step growth curves **(Fig. S2)**, are summarized in **Table 1**. For Φ*Xhv*-2, Φ*Xhv*-5, Φ*Xhv*-7, Φ*Xhv*-18 and Φ*Xhv*-28, the complete replication cycles (including adsorption) ranged from 95 min (Φ*Xhv*-7) to 135 min (Φ*Xhv*-5). Latent periods were short, ranging from 10 and 20 min, followed by a rise period lasting 40 to 80 min for phages Φ*Xhv*-28 and Φ*Xhv*-5, respectively. Burst sizes varied considerably, from 5 (Φ*Xhv*-28) to 31 (Φ*Xhv*-5) virions per infected cell. In contrast, Φ*Xhv*-12, Φ*Xhv*-15, and Φ*Xhv*-16, exhibited notably longer replication cycles (200-230 min), with latent periods approximately equal to the duration of the burst phase (∼1 h). Burst sizes were low for these phages, ranging from 2 (Φ*Xhv*-12) to 5 (Φ*Xhv*-15) virions per infected cell. These results were consistent with the infection curves and highlight notable difference in the lytic efficacy and dynamics of the newly phages. These characteristics, such as burst size and latent period, are crucial and should be carefully considered alongside host range and lytic activity when selecting phages for large-scale production for biocontrol applications.

**Figure 3.**
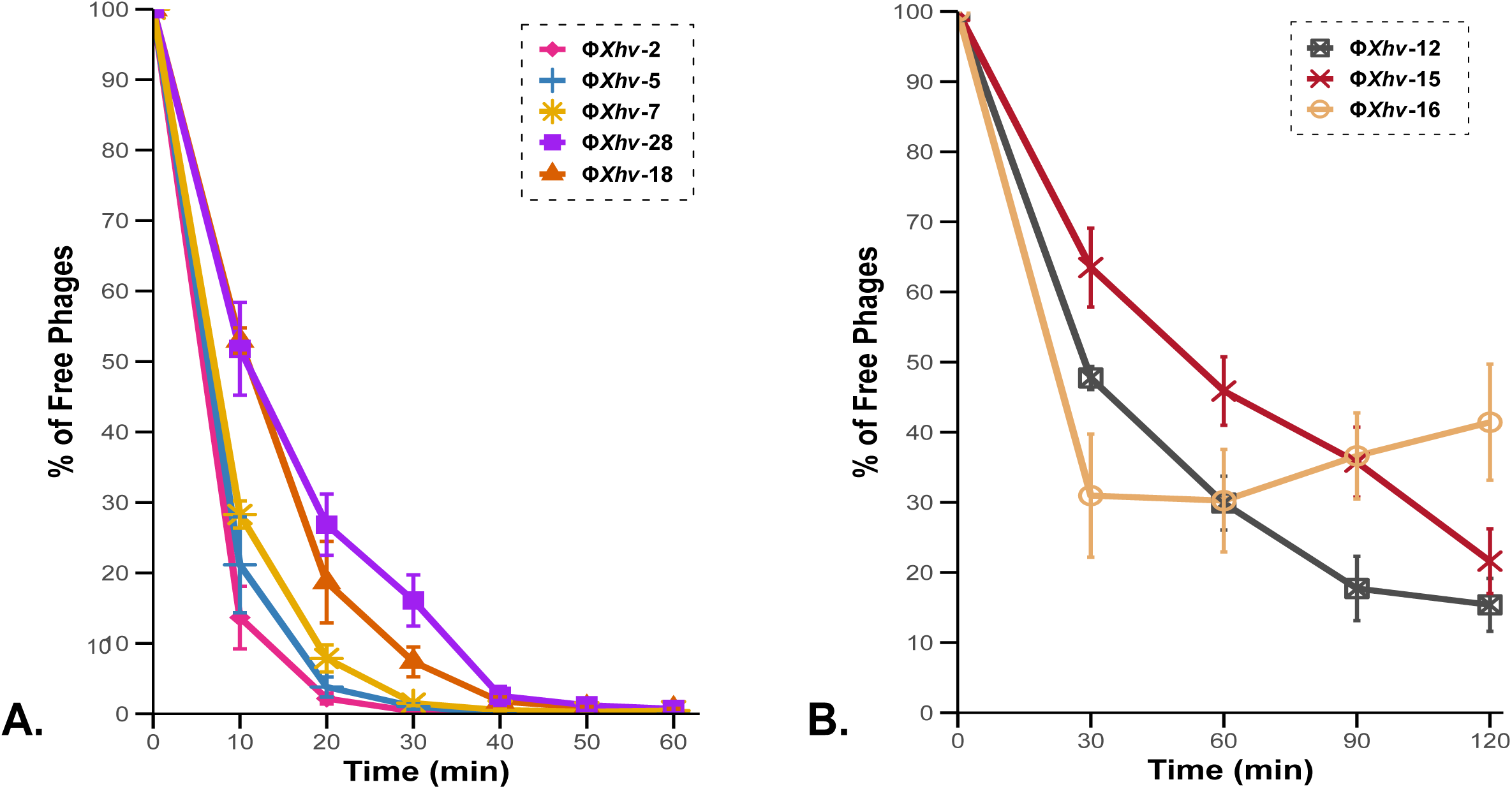
Adsorption kinetics of two distinct genomic groups of bacteriophages to *X. hortorum* pv. *vitians*. The adsorption efficiency of eight phages to their host was assessed by monitoring the percentage of free phages remaining in the supernatant over time. Assays were conducted at a MOI of 0.01 at 28°C. At each time point, aliquots were collected, filtered to separate free phages from adsorbed ones, and quantified using a standard spot assay. Data points represent mean ± standard error of the mean (SEM) from two independent biological replicates, each with three technical replicates. **A.** Adsorption curves of the smaller-genome phages: Φ*Xhv*-2, Φ*Xhv*-5, Φ*Xhv*-7, Φ*Xhv*-18, and Φ*Xhv*-28. These phages exhibited rapid adsorption kinetics, with over 90% of phages adsorbed within 20 to 30 min for most. **B.** Adsorption profiles of the three jumbophages (Φ*Xhv*-12, Φ*Xhv*-15, and Φ*Xhv*-16) exhibiting slower adsorption kinetics, characterized by a substantial proportion of free phages remaining unabsorbed even after 60 min.

**Table 1.** Biological and kinetic parameters of *Xanthomonas hortorum* pv. *vitians* phages

| Phage name | Adsorption rate<br>constant<br>K (mL.cell<sup>-1</sup>.min<sup>-1</sup>) | Latent period<br>(min) | Rise period<br>(min) | Lytic cycle duration<br>(min) | Burst size | Titer after<br>liquid culture (PFU/mL) |
| --- | --- | --- | --- | --- | --- | --- |
| ΦXhv-2 | $2.1 \times 10^{-7}$ | 10 | 70 | 115 | 7 | $2.93 \times 10^{10}$ |
| ΦXhv-5 | $1.3 \times 10^{-7}$ | 20 | 80 | 135 | 31 | $2.05 \times 10^{10}$ |
| ΦXhv-7 | $1.3 \times 10^{-7}$ | 10 | 50 | 95 | 28 | $2.85 \times 10^{10}$ |
| ΦXhv-18 | $3.8 \times 10^{-7}$ | 10 | 50 | 105 | 6 | $1.2 \times 10^{10}$ |
| ΦXhv-28 | $5.6 \times 10^{-7}$ | 20 | 40 | 115 | 5 | $8.93 \times 10^9$ |
| ΦXhv-12 | $7.5 \times 10^{-7}$ | 60 | 60 | 200 | 2 | $2.88 \times 10^9$ |
| ΦXhv-15 | $9.1 \times 10^{-7}$ | 60 | 60 | 230 | 5 | $3.51 \times 10^9$ |
| ΦXhv-16 | $5.7 \times 10^{-7}$ | 60 | 60 | 200 | 3 | $7.25 \times 10^9$ |

### Phages exhibit high stability under environmental stress but are sensitive to UV exposure

Phage stability was assessed under different physicochemical conditions, including variations in pH, temperature, and UV exposure (**Fig. 4**). All phages remained highly stable across a broad pH range (pH 6 to 10), with survival rates generally above 82%. Except for phages Φ*Xhv*-15 (at pH 12), as well as Φ*Xhv*-12, Φ*Xhv*-16, the newly isolated phages also survived in even more acidic (pH 4) and alkaline (pH 12) conditions, with moderate reductions in viability (less than 36%). However, survival dropped at pH 2 with near-complete or total inactivation (**Fig4.A**). Temperature tolerance profiles revealed that all phages remained highly viable at 28°C but were completely inactivated at 65°C and above, with survival rates falling below detection limits. At 28°C, survival ranged from 65% for Φ*Xhv*-16 to an average of 95.5% for the other phages. A reduction in phage viability was observed at 37°C for Φ*Xhv*-12, Φ*Xhv*-15, and Φ*Xhv*-16. At 55°C, these three phages were fully inactivated, while a slight decrease in survival was also detected for Φ*Xhv*-5 and Φ*Xhv*-18 (**Fig4.A**). UV exposure led to a rapid decrease in phage viability. After 2 min of exposure, a significant reduction was observed, with survival dropping close to zero between 5 and 10 min for all phages. Φ*Xhv*-2, Φ*Xhv*-5, Φ*Xhv*-18 and Φ*Xhv*-28 appeared slightly more UV-tolerant, retaining over 40% viability at 2 min, whereas Φ*Xhv*-12, Φ*Xhv*-15, and Φ*Xhv*-16 were more rapidly inactivated (**Fig4.B**). Overall, these results indicate that the phages are relatively stable under a range of conditions compatible with environmental applications, regarding pH and temperature. However, all phages showed marked sensitivity to UV exposure.

**Figure 4.**
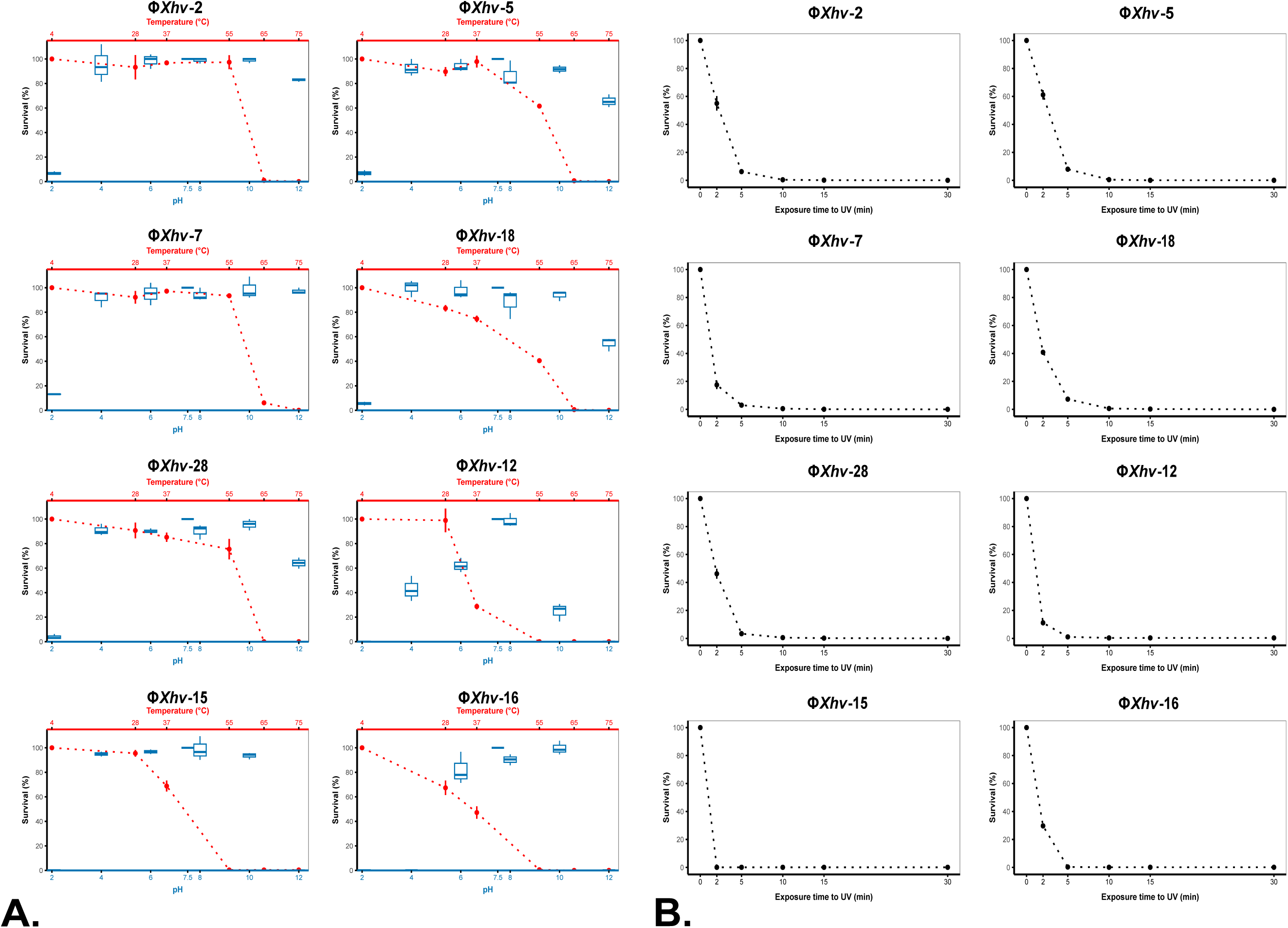
Abiotic stability of *X*. *hortorum* pv. *vitians* phages. The survival of eight bacteriophages was evaluated following exposure to various abiotic stress conditions, including temperature, pH, and UV radiation. In all assays, phage survival was quantified by spot assay and expressed as the percentage of infectious particles remaining relative to untreated controls (set as 100%, corresponding to initial phage titers at 4°C, in the dark, in SM Buffer at pH 7.5). Values represent mean ± standard error of the mean (SEM) of two independent biological replicates, each with three technical replicates. **A.** Thermal and pH stability. Each panel corresponds to an individual phage’s survival profile. Temperature stability (red dotted line) was evaluated after 1-hour incubation in the dark at temperatures ranging from 4°C to 75°C (with temperature values indicated on the upper x-axis). pH stability (blue box plots) was determined after1-hour incubation at room temperature in the dark, in SM Buffer across a pH gradient ranging from 2 to 12 (pH values indicated on the lower x-axis). **B.** UV sensitivity. Phage survival following exposure to UV-B radiation (312 nm) was monitored over a 30-minute period. Survival rapidly declined with increasing exposure time.

### Genomic features highlight two distinct phage lineages

Whole-genome sequencing analysis of the novel *X*. *hortorum* pv. *vitians* phages are summarized in **Table 2**. All phage genomes were linear, double-stranded DNA molecules. Based on their similar genome sizes, the phages were grouped into two categories: the smaller-genome phages Φ*Xhv*-2, Φ*Xhv*-5, Φ*Xhv*-7, Φ*Xhv*-18 and Φ*Xhv*-28 (46,786 - 47,861 bp), and the jumbophages Φ*Xhv*-12, Φ*Xhv*-15, Φ*Xhv*-16 (227,648 - 229,124 bp). The averages of GC contents were 61.8% and 55.1% for the two groups, respectively. The small-genome phages encoded between 72 and 76 predicted CDSs, with over half annotated as hypothetical or unknown proteins. In contrast, the jumbophages harbored between 238 and 330 CDSs, of which more than 75% encode proteins of unknown function (**Table S3**). Functionally categorized proteins were classified into 5 categories: replication-related, structural and morphogenesis, lysis and host interaction, hypothetical and others proteins (**Fig. 5**). A single tRNA gene encoding asparagine (Asn), with the anticodon GTT, was identified in the jumbophage genomes. The preferred Asn codon in both phages and their bacterial host was AAC, corresponding to this anticodon. Differential codon usage analysis revealed that AAC was the second most overrepresented, with a usage frequency 1.5 times higher in phages Φ*Xhv*-15 and Φ*Xhv*-16 than in the host (**Fig. S3**). On average, 59.56% of CDSs in the phage genomes either exclusively used AAC for Asn or showed a bias toward AAC over the codon AAT. Among annotated genes, those with strongest AAC bias were mostly associated with structural and morphogenesis roles, including tail proteins, the major head protein, baseplate hub subunits, head maturation proteases, and minor tail proteins (**Table S4**). No antibiotic resistance genes, virulence factors, or toxins were detected in the genomes based on analyses using the CARD, ResFinder, and VFDB databases. A virulent lifestyle was predicted for all phages, which is consistent with the absence of lysogeny-related genes, such as integrases. Together, these genomic features support the potential suitability of these phages for biocontrol applications.

**Table 2.** Genomic features of eight novel *Xanthomonas hortorum* pv. *vitians* phages

| Phage Name | Accession Number | Reference Host | Genome Size (bp) | GC Content (%) | No. of CDSa | % of known functions | No. of tRNAa | Lifestyle Predictionb |
| --- | --- | --- | --- | --- | --- | --- | --- | --- |
| ΦXhv-2 | PV702347 | CFBP8642 | 47,229 | 61.88 | 74 | 51 | 0 | virulent (99.66%) |
| ΦXhv-5 | PV764598 | CFBP8639 | 47,720 | 61.64 | 76 | 46 | 0 | virulent (99.95%) |
| ΦXhv-7 | PV764599 | CFBP8641 | 46,786 | 62.05 | 72 | 36 | 0 | virulent (99.91%) |
| ΦXhv-18 | PV764600 | CFBP8643 | 47,861 | 61.60 | 73 | 49 | 0 | virulent (99.60%) |
| ΦXhv-28 | PV754127 | AB16734ΔLPS3 | 46,883 | 61.86 | 72 | 51 | 0 | virulent (97.64%) |
| ΦXhv-12 | PV764601 | CFBP8640 | 227,648 | 55.11 | 330 | 24 | 1 | virulent (100%) |
| ΦXhv-15 | PV764602 | CFBP8640 | 228,725 | 55.17 | 328 | 23 | 1 | virulent (100%) |
| ΦXhv-16 | PV764603 | CFBP8640 | 229,124 | 55.09 | 330 | 23 | 1 | virulent (100%) |

<sup>a</sup>Structural and functional annotation was carried out using the rTOOLS pipeline (RIME Bioinformatics). Coding sequences (CDS) and tRNAs were predicted using Prodigal v2.6.3, Phanotate v1.5.0, Glimmer v3.02, GeneMarkS v1.14, and tRNAscan-SE v2.0.7. Functional annotation of all predicted CDS was performed using a combination of databases: CARD (2023.11), ResFinder (2023.04), VFDB (2023.11), Phantome (2021.03), Swiss-Prot (2023.11), NCBI\_virus (2023.11), VOG (2020.05), pVOG (2021.03), RefSeq (2023.11), and PHROGs (2023.02). <sup>b</sup>Phage lifestyle was predicted by the machine-learning-based program PhageAI and, further, confirmed by the absence of integrase genes.

**Figure 5.**
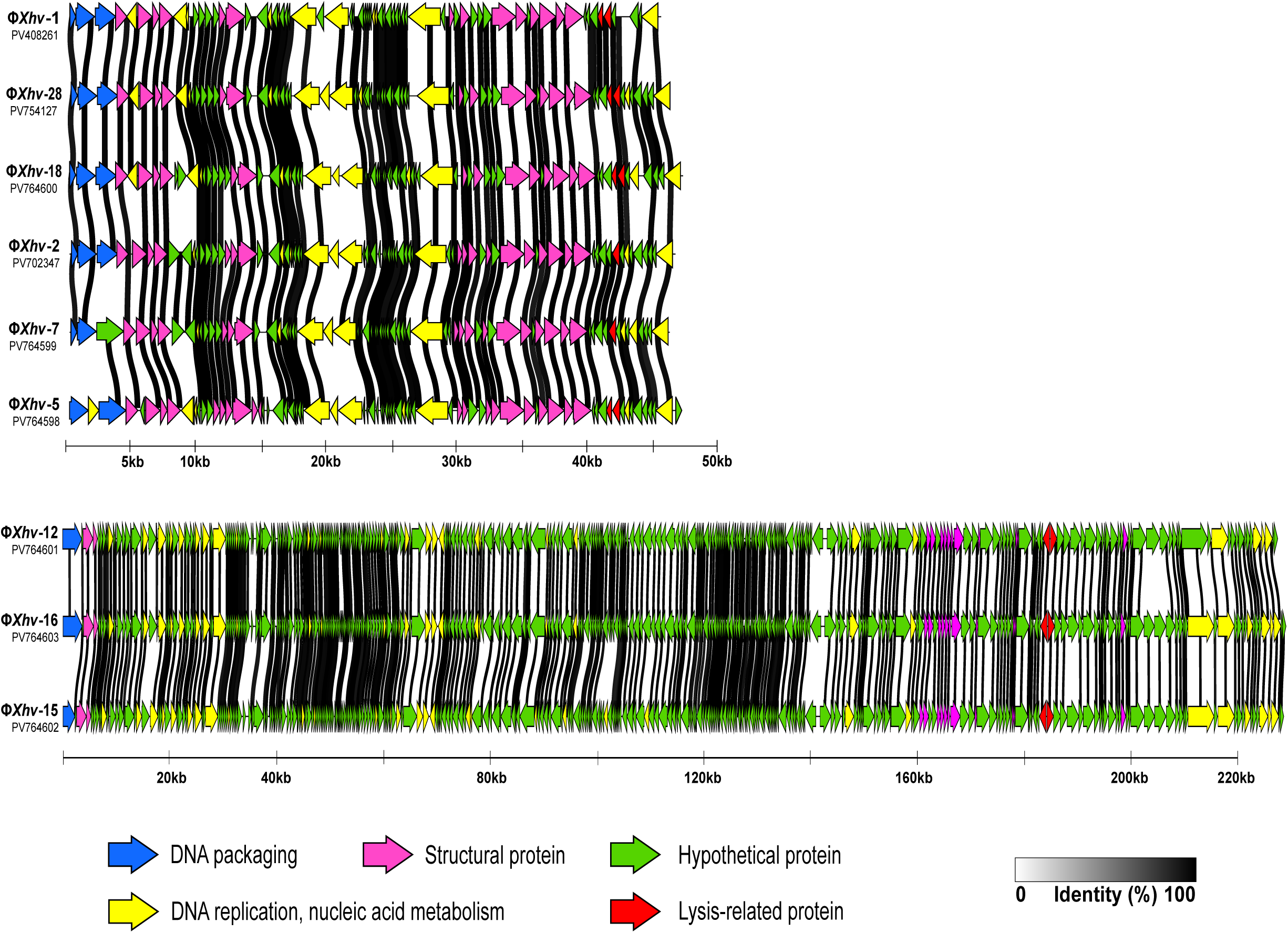
Comparative genomic organization of *X. hortorum* pv. *vitians* bacteriophages genomes. The figure displays the genomic organization, syntheny, and sequence similarity among bacteriophages infecting *X*. *hortorum* pv. *vitians*. Phages are grouped by genome size into smaller-genome phages (top panel) and jumbophages (bottom panel). Arrows represent predicted open reading frames (ORFs) and indicate transcriptional orientation. ORFs are color-coded by predicted function: DNA packaging (blue), structural protein (magenta), DNA replication and nucleic acid metabolism (yellow), lysis-related protein (red), and hypothetical protein (green). Shaded connectors between genomes show pairwise amino acid sequence identity, with gradient intensity representing identity from 0% (white) to 100% (black); only identity ≥ 90% are displayed. GenBank accession numbers for each phage genome are provided on the left. Genome comparisons were generated using Clinker and edited in Inkscape v1.4.

### Comparative genomics confirm lineage divergence and suggest a new jumbophage genus

To provide an overview of the phylogenetic relationships of the newly isolated phages with other phages, a whole-proteome-based phylogenetic tree was generated with ViPTree using reference genomes from the Virus-Host DB (45) (**Fig. S4**). The phages clustered into two major groups, consistent with their genome sizes and divergence in biological properties. The jumbophages were related to phages infecting various bacterial hosts, such as *Pseudomonadota* (*Ralstonia*, *Burkholderia*, *Pseudomonas*, *Dickeya*), *Actinomycetota* (*Propionibacterium*), and *Bacillota* (*Clostridium*). In contrast, the smaller-genome phages clustered more closely with other *Xanthomonas* phages and other *Pseudomonadota*-associated phages. These observations were further refined by VIRIDIC analysis, which quantified the intergenomic similarities between the eight novel phages and closely related genomes available in GenBank (**Fig. 6**). The smaller-genome phages shared a high percentage of intergenomic similarity (> 82%) with each other and with the previously described phage Φ*Xhv*-1 (PV408261), a lytic phage infecting *X*. *hortorum* pv. *vitians* LM16734, forming a coherent genus. These phages also displayed moderate similarity (i.e., 45 to 50.5 %) with lytic *Xanthomonas* sp. phages BsXeu269p/3 (ON996340.1), KPhi1 (NC_054460.1) and MYK3 (OK275494.1). Although these phages belong to a distinct phage genus according to the ITCV guidelines, their genomic relatedness suggests a common evolutionary lineage. This genomic proximity, combined with divergence in sequence, may contribute to explain their narrow host range strictly limited to pathovar *vitians* strains. In contrast, the jumbophages exhibited high similarity within their group (> 95%), consistent with their clustering in the proteome-based three. However, they shared no detectable identity with the five smaller-genome phages or with any other phages in the database. According to ICTV thresholds, this supports their classification as members of a novel phage genus, tentatively named *Xanthorvitianjumbovirus*, with *Xanthorvitianjumbovirus xhv12like* as the proposed species name. Whole-genome synteny analysis using Clinker revealed conserved modules within each of the two phage groups (**Fig. 5**), further supporting the genomic coherence of the distinct phage lineages. The jumbophage genomes (Φ*Xhv*-12, Φ*Xhv*-15, and Φ*Xhv*-16) displayed near-perfect synteny and high nucleotide identity, consistent with their classification as members of the same phage species. This genomic similarity may also underlie their similar infection profiles. In summary, these results reveal two distinct phage lineages, small-genome phages and jumbophages, which differ in their infection dynamics, host range, biological properties, and genomic architecture.

**Figure 6.**
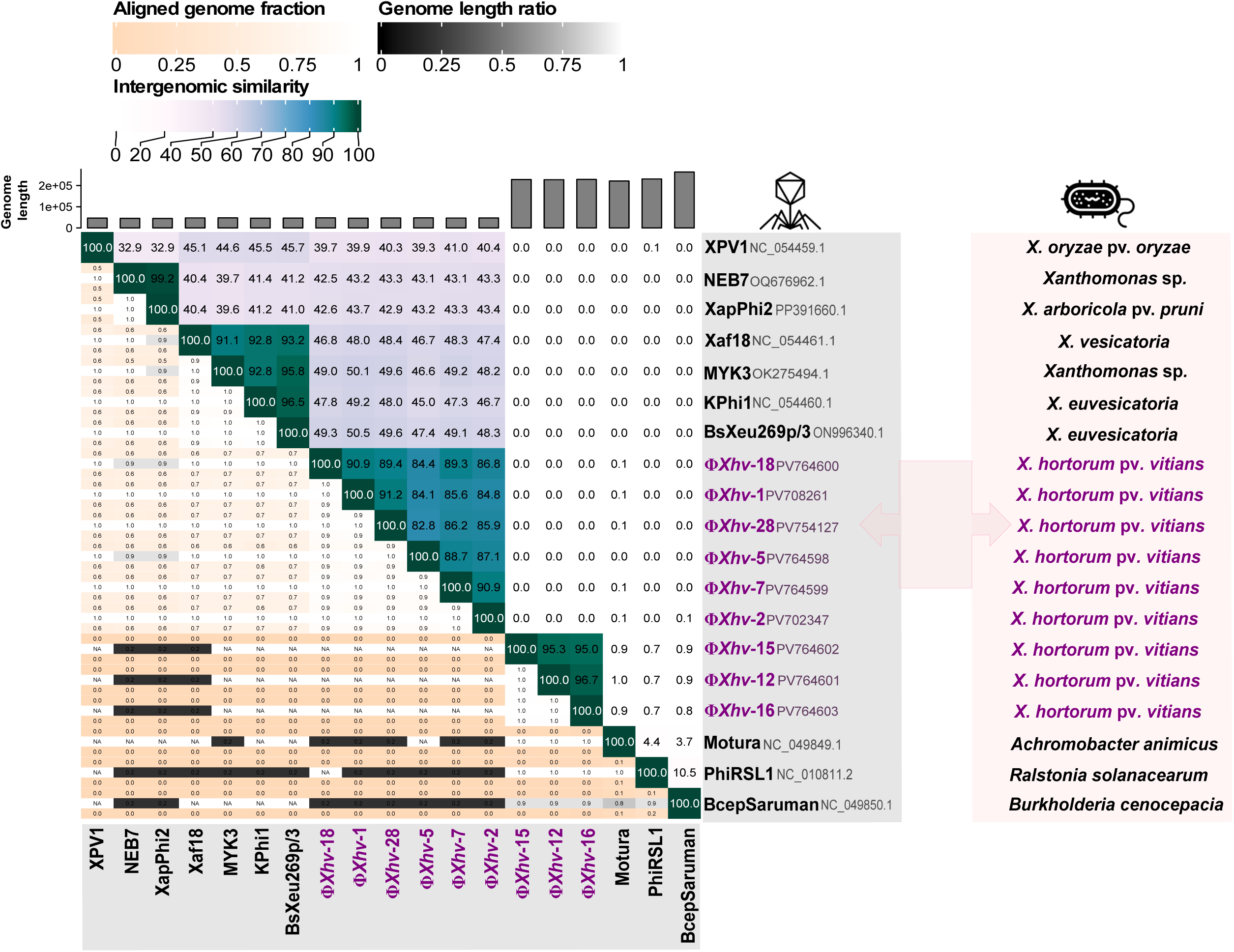
Intergenomic similarity analysis of *X*. *hortorum* pv. *vitians* phages and related bacteriophages. Pairwise genome comparisons between the novel *X*. *hortorum* pv. *vitians* phages and their closest known relatives were generated with VIRIDIC and edited using Inkscape v1.4. The matrix displays comparative genomic metrics for each phage pair. The upper-right triangle of the matrix (white to green color scale) represents intergenomic similarity (%), with darker green indicating higher similarity. The lower-left triangle (peach scale) shows the aligned fraction of each genome pair. The bar chart above the matrix displays the genome length of each phage. The right panel lists the predicted bacterial host of each phage. Phages are labeled with their names and corresponding GenBank accession numbers.

## DISCUSSION

The isolation and characterization of bacteriophages are critical steps for developing effective biocontrol strategies against phytopathogenic bacteria. To date, 195 complete genome sequences of phages infecting *Xanthomonas* have been reported, of which only nine are classified as jumbophages (GenBank database, last accessed 2 August 2025). In this study, we report on eight newly isolated phages active against *X*. *hortorum* pv. *vitians*, including 3 jumbophages, the first known to infect this pathovar. Through detailed analysis of their host range, lytic cycle parameters, genetic features, and stability under physicochemical stress conditions, this study provides a comprehensive framework to assess the potential of these phages for biocontrol applications.

The jumbophages Φ*Xhv*-12, Φ*Xhv*-15, and Φ*Xhv*-16 were classified into a new lineage due to the complete absence of genome homology with phages available in public databases. Jumbophages remain underrepresented in current databases, accounting for fewer than 1% of sequenced phages, which may partly explain the lack of close genomic relatives for these isolates (46, 47). This underrepresentation primarily stems from the technical challenges inherent in their isolation, notably limited diffusion though standard agar concentration and physical retention during typical filtration steps (48). These inherent difficulties underscore the critical importance of adapting standard isolation protocols to ensure the effective detection and characterization of jumbophages, as exemplified by strategies employing lower agar concentrations or larger-pore filters for improved recovery.

These three new jumbophages share biological characteristics similar to those of jumbophages infecting phytopathogenic bacteria, although it remains unclear whether these traits reflect specific adaptations associated with their genome size. For instance, the phage vB_XciM_LucasX, which infects *Xanthomonas citri* and *X. fuscans*, has been reported to exhibit similar survival characteristics under abiotic stresses: (i) stability across pH ranges, with inactivation at pH 3 and 11; (ii) significant UV sensitivity, with only 25% survival observed after 80 s of exposure; and (iii) inactivation at temperatures above 50°C (49). Other jumbophages, such as Xac1 and Xbc2 which infect *Xanthomonas vesicatoria*, as well as RsoM2, infecting *Ralstonia solanacearum*, have been found to be more thermotolerant (50, 51). Compared to the small-genome phages isolated in this study, most jumbophages described to date, including Φ*Xhv*-12, Φ*Xhv*-15, and Φ*Xhv*-16, have demonstrated broader host spectra (52–54). Indeed, Φ*Xhv*-12, Φ*Xhv*-15, and Φ*Xhv*-16 lysed 74% of tested *X*. *hortorum* pv. *vitians* strains, and other jumbophages such as RsoM2, vB_XciM_LucasX, and XacN1 (55) have been reported to lyse on average more than 90% of their respective target hosts. As highlighted by a recent large-scale study in *E*. *coli*, host range is influenced by the genetic and structural diversity within the target bacterial species, particularly through polymorphisms in surface receptors (56). However, the broader host spectra observed here among jumbophages, compared to smaller-genome phages targeting the same species, suggest an additional contributing factor: a reduced dependence on host cellular machinery for genetic replication. This hypothesis is supported by the enriched gene content typically found in jumbo genomes. Jumbophages have been shown to encode a wider array of genes related to DNA replication, nucleotide metabolism, cell wall degradation (e.g., endolysine, glycoside hydrolase, chitinase), and tRNAs (47, 57, 58). In the present study, the jumbophages contained a greater number of replication-associated genes, including three DNA polymerases, four helicases, and five endonucleases, compared to only one of each gene type in the smaller-genome phages. In contrast to previously characterized jumbophages, which typically harbor around 15 tRNAs, Φ*Xhv*-12, Φ*Xhv*-15, and Φ*Xhv*-16 each encoded only one tRNA gene. Phage-encoded tRNAs are thought to play multiple roles. Several studies have proposed that phage-encoded tRNAs compensate for differences in codon usage by supplementing codons more frequently utilized by the phage than by the host (59). Other studies have also demonstrated that they can serve as a mechanism to evade host antiphage defense systems, such as retrons and tRNA-targeting nucleases, thereby facilitating successful infection and phage propagation (60, 61). In Φ*Xhv*-12, Φ*Xhv*-15, and Φ*Xhv*-16, the single tRNA identified, targeting the AAC codon for asparagine, corresponds to a notable codon usage bias observed in these phages and is predominantly associated with structural genes. While this suggests some degree of translational optimization, the limited tRNA repertoire of these phages may partly explain the slower infection kinetics with low burst sizes observed, as their translation efficiency remains more constrained compared to jumbophages that possess a more extensive set of tRNA. Consistently, the infection dynamics of these three jumbophages were characterized by long replication cycles, comparable to those reported for other jumbophages such as vb_XciM_LucasX (50, 51) and low burst sizes ranging from only 2 to 5 virions per infected cell. These values are among the lowest ever reported for phages infecting *Xanthomonas* (62, 63), and are comparable to those observed in other *myovirus* jumbophages such as Deimos-Minion and RAY, which infect *Erwinia* sp. (∼5 virions per infected cell) (64) or KTN4 (∼6-8 virions) infecting *Pseudomonas aeruginosa* (65). Other factors may also contribute to their slow replication dynamics. For instance, although a nucleus-like structure has not yet been described for jumbophages infecting phytopathogenic hosts, such compartments have been described in other jumbo *myoviruses* such as KTN4. This proteinaceous structure protects the phage genome from the bacterial defensome while imposing a fitness cost on replication efficiency (66). A similar mechanism could potentially operate in the jumbophages of this study, which might explain their observed low burst sizes.

In contrast to jumbophages, the smaller-genome phages isolated in this study exhibited biological and kinetics parameters consistent with *Xanthomonas* phages reported by Nakayinga collaborators (62). Indeed, most of the phages described in their review exhibited short latent periods ranging from 20 to 45 min and burst sizes varying from 4.6 to 350 virions per infected cell. Similar characteristics were observed for the five new phages characterized in our study, which demonstrated even shorter latent periods of 10 to 20 min and burst sizes ranging from 5 to 31 virions per cell. However, their high survival rates at pH values above 10 are uncommon but have also been reported for the phage BsXeu269p/3, which is active against *X. vesicatoria* and represents the closest genetic relative of the phages isolated in this study (67). A second phage, Kф1, which is also active against *X. vesicatoria* and shares over 40% of intergenomic similarity with the phages isolated in this study, has been reported to exhibit higher UV resistance and thermal tolerance up to 70°C, compared to the maximum survival temperature of 55°C observed in phages infecting *vitians* (68).

Although regulatory frameworks governing the use of bacteriophages as biocontrol agents remain under development and vary between countries, several critical criteria have been proposed to ensure the safe and effective use of phages in agriculture (69). From a genetic perspective, phages selected for biocontrol applications should be strictly lytic, as confirmed by the absence of lysogeny-associated genes, such as integrases or site-specific recombinases. In addition, these phages must not carry any known toxin or antibiotic resistance genes (9). Ideally, they should also encode endonucleases to minimize their role as “superspreaders” by degrading bacterial DNA released during infection, thereby reducing the risk of horizontal gene transfer through natural transformation (70). These genetic requirements are fully met by the newly isolated phages described in this study. No genes associated with lysogeny, toxins, or antibiotic resistance were detected, consistent with a strictly lytic lifestyle. Furthermore, nearly all phages encode at least one endonuclease, with up to three endonucleases identified in the genomes of the jumbophages.

From successful biocontrol, phages must retain biological activity and stability in environmental conditions (15). Tailed phages in particular are known to exhibit higher stability when exposed toadverse environmental conditions (71, 72). In this study, the phages demonstrated thermal stability up to 55°C and maintained infectivity across a broad pH range of 4 to 10. However, they were rapidly inactivated by UV exposure, with complete loss of activity occurring within 5 min. Since *X*. *hortorum* pv. *vitians* is a foliar pathogen, improving phage persistence on leaf surfaces in crucial for ensuring their efficacy in field conditions (17). Various strategies have been explored to address this challenge, including late-evening applications (73), co-application with non-pathogenic carrier bacteria or plant extracts, and protective formulations to mitigate UV inactivation (74–76). Among these, Balogh *et al*. (77) showed that a formulation containing 0.5% Casecrete NH-400 (a water-soluble casein protein polymer), 0.5% sucrose, and 0.25% pregelatinized corn flour increased phage survival on the tomato phyllosphere by approximately 1,000-fold compared to unformulated phages. Similarly, the addition of skim milk and sucrose improved phage persistence and contributed to disease reduction in citrus canker (75, 78). More recently, innovative strategies such as incorporating a food-grade dye (brilliant blue FCF) or encapsulation in nanoparticules (e.g., NAC-ZnS or chitosan-coated Ca-alginate nanoparticles) have been reported to protect phages from UV inactivation (79–81).

While such formulations have been developed to enhance phage persistence on the phyllosphere, some genetically related phages to those isolated in this study, such as BsXeu269p/3 and Kф1, have already demonstrated promising biocontrol efficacy without requiring protective formulations. For instance, BsXeu269p/3 effectively reduced the spread of *X*. *euvesicatoria* in pepper plants by limiting pathogen transmission via aerosols and decreasing bacterial loads in local lesions (82). Similarly, Kф1 persisted on pepper leaves for up to 7 days under greenhouse conditions, and significantly reduced disease severity with dual applications administered before and after pathogen inoculation (83).

Finally, effective phage biocontrol requires both high infectivity and specificity against the target pathogen (9). The complementary infection profiles of our newly isolated phages highlight the potential of phage cocktails to broaden the host range while maintaining specificity to the pathovar. The design of phage cocktails must also address the potential emergence of bacterial resistance, a major challenge in phage-based biocontrol (84). One promising approach involved targeted isolation of phages against resistant bacterial mutants (85), or the use of evolved phages exhibiting expanded host ranges (86). Consistent with this strategy, Φ*Xhv*-28 was isolated using a bacterial mutant resistant to Φ*Xhv*-1. Resistance in this mutant is mediated by mutations in the O-antigen side chains, the primary receptor of Φ*Xhv*-1. Although Φ*Xhv*-28 has a narrow host range restricted to this resistant mutant, its ability to specifically recognize altered receptor structures underscores its potential to complement broader-spectrum phage cocktails, supporting robust biocontrol of *X*. *hortorum* pv. *vitians*.

## CONCLUSION

Despite the recognized potential of bacteriophages in agricultural biocontrol, their diversity and utility against many plant pathogens remain largely underexplored. This study characterizes a collection of eight novel phages, including the first jumbophages identified as active against *Xanthomonas hortorum* pv. *vitians*. Distinct biological and genomic profiles have been revealed among these phages. The newly identified jumbophages, classified into a novel lineage, exhibit broad host ranges and low burst sizes, which may be linked to their minimal tRNA repertoire and possible formation of intracellular compartmentalization. In contrast, smaller genome phages demonstrate classic lytic cycles with shorter latent periods and higher burst sizes. Both groups have exhibited good thermal and pH stability compatible with foliar field application, though all phages were highly sensitive to UV irradiation. Crucially, the genetic profiles of all isolated phages fully comply with established biocontrol criteria, confirming their strictly lytic nature and absence of lysogeny-associated, toxin, or antibiotic resistance genes, while encoding endonucleases. The complementary biological features of our jumbophages and smaller-genome phages, especially their diverse host ranges and burst sizes, highlight their combined potential in tailored phage cocktails for robust management of *X*. *hortorum* pv. *vitians* infections. To fully harness this potential, further studies are warranted to characterize phage-host interactions at the molecular level, including detailed receptor identification, which is crucial to anticipate and mitigate the emergence of bacterial resistance. In parallel, formulation strategies should be investigated to enhance phage persistence under UV exposure, a known limitation for many phages. Rigorous efficacy trials in greenhouse and field settings will also be essential to validate the practical effectiveness of these phages for crop disease management. Overall, this study significantly expands our understanding of bacteriophages infecting *X*. *hortorum* pv. *vitians,* providing a critical foundation for developing effective, phage-based solutions against this important foliar pathogen.

## Author contributions

A.B conceptualized the study, conducted the investigations, curated and analyzed the data, created the visualizations, and wrote the original draft. I.R participated in all experimental work and contributed to data analysis. N.T was involved in phage isolation, purification, and production. M.G coordinated R&D activities at GREENPHAGE, contributed to experimental design, ensured the quality and reproducibility of results, and performed TEM imaging. D.C contributed to methodology development and provided specific resources. F.B supervised the manuscript, acquired funding and managed the overall project. All authors reviewed and edited the manuscript.

## Data availability statement

The assembled and annotated genomes of the eight novel phages described in this study have been deposited in GenBank under accession numbers PV702347, PV64598, and PV764599 to PV764603. All datasets supporting the conclusions of this article are available within the article and its Supplementary Information files. In addition, raw experimental data from wet lab assays have been made publicly available via Zenodo (DOI:<u>10.5281/zenodo.16728738</u>) (1). During the peer review process, data remain under restricted access; reviewers may access them directly via the following private link: https://zenodo.org/records/16728739?token=eyJhbGciOiJIUzUxMiJ9.eyJpZCI6IjUxNDk2MDQ0LTMwNWMtNGVmMy05ODMwLTkyOTA4MjRhNDc0ZiIsImRhdGEiOnt9LCJyYW5kb20iOiJkNGMzYmMyNDBjY2FlNWExNWU2NTMxOWU2NzhlZmFiOSJ9.swpqi6c4JiiaW4_Anpv0DYkyKx4DaK6MSK8varlyh7N-nkxusx-SpuXMjMuivlzagC6ilOz8RpwIAmzXLPGxSw)

## Acknowledgments

The authors are grateful to the FNX and Phages.fr networks for their insightful discussions during scientific meetings. We acknowledge Concepcion Sanchez-Cid Torres for preparing the library and performing the sequencing run of phage Φ*Xhv*-7. We also thank Danis Abrouk for his preliminary sequence analyses of Φ*Xhv*-7 and for his valuable advice on statistical analyses. Finally, we warmly thank Laurène Robert and Hanane Amari for their valuable contribution to the isolation and characterization of certain phages included in this study.

---

**Figure S1. Growth kinetics of *X*. *hortorum* pv. *vitians* after infection by phages at different multiplicities of infection (MOIs).** Each panel displays the mean bacterial growth curve measured by optical density at 600nm (OD_600nm_) over 24 h at 28°C for a specific phage (Φ*Xhv*-2, Φ*Xhv*-5, Φ*Xhv*-7, Φ*Xhv*-12, Φ*Xhv*-15, Φ*Xhv*-16, Φ*Xhv*-18, or Φ*Xhv*-28). Phages were added at MOIs ranging from 0.00001 to 50, as indicated by the color-coded legend. Phage-free controls (0 MOI, Wild Type -WT) are shown in grey. Shaded areas represent the standard deviation from four technical replicates per condition.

**Figure S2. One-step growth curves of phages infecting *X*. *hortorum* pv. *vitians*.** Each panel shows the phage titer (PFU/mL) measured over time following infection of exponentially growing *X*. *hortorum* pv. *vitians* cultures with individual phages: Φ*Xhv*-2, Φ*Xhv*-5, Φ*Xhv*-7, Φ*Xhv*-12, Φ*Xhv*-15, Φ*Xhv*-16, Φ*Xhv*-18, and Φ*Xhv*-28. Bacterial cells were infected at a MOI of 0.1 and incubated at 28°C. Samples were collected at regular intervals, and the number of infection particles released was determined by spot assay. Dashed lines indicate the assay detection limit. Data points represent mean ± standard deviation from two independent biological replicates, each with three technical replicates per time point.

**Figure S3. Codon usage bias of jumbophages relative to *X*. *hortorum* pv. *vitians*.** The bar plot illustrates the difference in codon usage frequency (Δ frequency = phage - host) for each of the 64 DNA codon, comparing the three jumbophages (Φ*Xhv*-12, Φ*Xhv*-15, and Φ*Xhv*-16) and their host, *X*. *hortorum* pv. *vitians* strain CFBP8640. Positive values indicate codons more frequently used by the phage than the host, whereas negative values indicate codons less frequently used. Codons are ordered alphabetically along the x-axis. The red star highlights codon “AAC” (Asn), which corresponds to the tRNA-Asn (anticodon GTT) encoded by the jumbophages.

**Figure S4. Phylogenetic analysis of *X. hortorum* pv. *vitians* bacteriophages within a global context of phage diversity.** This proteomic tree depicts the phylogenetic positions of eight newly discovered *X*. *hortorum* pv. *vitians* phage genomes (indicated by red stars) relative to 5,640 reference phage genomes. The tree was generated with VipTree and edited using Inkscape v1.4. The inner ring indicates virus family assignments, while the outer ring shows host group classifications of these phages. Two zoomed-in subtrees highlight the phylogenetic positions of the *X*. *hortorum* pv. *vitians* phages relative to closely related viruses. Branch lengths represent genome-wide proteomic distances. Phages are labeled with their respective GenBank accession numbers.

**Table S1.** List of *Xanthomonas* strains used in this study.

**Table S2.** Morphological features of eight novel *Xanthomonas hortorum* pv. *vitians* phages.

**Table S3.** Predicted coding sequences and functional annotations of phage genomes.

**Table S4.** Asparagine codon bias in coding sequences of *Xanthomonas hortorum* pv. *Vitians* jumbophages.

